# A 50-marker mass cytometry panel to expand analysis of the functional breadth of human immune cells

**DOI:** 10.64898/2026.06.03.729939

**Authors:** Laura C. Polanco, Michael J. Cohen, Lauren Tracey, Christina Loh, Erika L. Smith-Mahoney, Amedeo J. Cappione, David King, Anna C. Belkina, Jennifer E. Snyder-Cappione

## Abstract

Human immune single-cell proteomic functional profiling has historically been performed with a limited number of inflammatory and/or cytotoxic readouts, capturing only a fraction of the complex orchestra of factors that comprise immune responses. Given the rising global crisis of chronic inflammation and the lack of clinically available treatment options, there is an urgent need to gain insight into the cell subsets that exhibit anti-inflammatory functional profiles and elucidate the mechanisms regulating these effector capacities. To address this, we developed a 50-marker CyTOF panel that enables unprecedented functional fingerprinting of human T cells, NK cells, monocytes, and B cells, detecting 24 intracellular targets. Healthy donor PBMCs were stimulated *ex vivo* and stained with this panel; from T cells, cytokines associated with the hallmark Type 1 (IFN-γ, TNF-α), Type 2 (IL-4, IL-5, and IL-13), and Type 17 (IL-17A, IL-17F) functional lineages were detected, as well as the chemokines MIP-1-α, MIP1-β, and IL-8 and the cell repair factor amphiregulin; from monocytes, IL-1β, IL-35, and IL-8 were detected. To ascertain if some of the cytokines less commonly included in Intracellular Cytokine Staining (ICS) panels were produced in response to physiological TCR stimulation via viral peptides, we measured the T cell response to a CMV-EBV-Flu (CEF) pool; in addition to TNF-α, IFN-γ, and IL-2, we also found that individual T cells produced additional cytokines with IFN-γ and TNF-α, such as amphiregulin, MIP-1α, IL-13, and IL-4. This mass cytometry panel provides an exceptionally broad and deeply resolved view of the functional diversity of human immune cells, surpassing, to our knowledge, the capabilities of previously reported approaches. Due to minimal signal overlap, CyTOF enables flexible panel customization, allowing markers and metal tags to be readily expanded or modified. Based on its resolution and adaptability, we anticipate that this panel and its derivatives will enable the discovery of novel immunomodulatory mechanisms for therapeutic intervention.

## Introduction

Immune responses are immensely complex, with intricate exchanges of chemokines, cytokines, and other mediators that collectively orchestrate inflammation, cell recruitment, cell death, and quiescent return to tissue homeostasis once the response has ended. Disease-associated immune responses involve heterogeneous combinations of functional states; resolving diverse functions within individual cells is critical for understanding immune regulation, cellular interplay, and clinical outcomes. For multiple reasons, including a shift toward multicellular and higher-parameter analysis and the difficulty in resolving many cytokines, such as IL-5, IL-10, and IL-13, by fluorescent cytometry, the discovery of key facets of immune responses beyond Th1 inflammation/cytotoxicity has progressed very slowly for decades. Our limited understanding of the full spectrum of functional proteins elicited in response to immune cell activation is set against the backdrop of a global epidemic of chronic inflammation, in which blocking inflammatory mediators or steroids has been the standard of care, with limited effective long-term solutions.^1^

To develop efficacious therapeutic interventions for many illnesses driven, in whole or in part, by chronic inflammation, including cancer, autoimmunity, and neurodegenerative diseases, detection of the full panoply of effector functions elicited from individual cells during immune responses is required. In addition, despite great success in CAR T therapy for B-cell cancers and the immense promise of cellular therapies to treat a variety of conditions with currently limited treatment options, overall progress in this area has stalled. This is likely due, at least in part, to what we predict is an underestimation of the functional breadth within products, including functions that directly counter the therapeutic activity of the drug, with such counter/opposing functions possibly exerted simultaneously with the desired activity from the same cell.

## Panel Rationale and Results

Many cytokines have historically been challenging to detect in human cells using fluorescent cytometry; as a result, orthogonal methods, such as the Enzyme-Linked Immunospot (ELISPOT) assay, have been used, enabling high-sensitivity detection of chemokines and Th2 cytokines.^2^ However, ELISPOT assays do not allow the simultaneous multiplexing of intracellular and surface marker expression of individual cells. Therefore, our group conducted a direct head-to-head comparison of fluorescent and mass cytometry for the intracellular detection of cytokines, transcription factors, and phosphoproteins. We reported that while some readouts, such as IFN-γ and IL-4, have similar resolution between the two platforms, Cytometry by Time of Flight (CyTOF) was overall clearly superior for detection of intracellular targets, with some readouts not detected clearly by fluorescent panels, despite high frequencies of cytokine-producing cells in the sample (i.e., IL-13).^3^

To further elucidate the functional and phenotypic diversity of human immune cells after short-term stimulation (*ex vivo*) from human blood (PBMC) and other biological specimens, and to functionally and phenotypically fingerprint cell therapy products, we designed and optimized a 50-parameter mass cytometry panel containing 24 intracellular targets. To our knowledge, this is the largest cytometry panel to date developed for multiplex profiling of immune cells spanning many major functional lineages (e.g., Th1, Th2, Th17, Treg). The panel evaluates functional signatures of a variety of circulating immune cell populations, such as αβ (adaptive) T cells, unconventional T cells (including mucosal-associated invariant T (MAIT) cells, invariant natural killer T (iNKT) cells, and γδ T cells), NK cells, B cells, and myeloid cells/monocytes. The effector functions measured span across major T cell linages, as well as chemokines and innate inflammatory and cell-repair cytokines. With this panel, we robustly detected cytokine signals following overnight stimulation of cryopreserved PBMCs from healthy donors. In addition, overnight stimulation with viral peptides derived from CMV, EBV, and influenza virus (‘CEF’ pool) revealed, to our knowledge, previously unreported cytokines and chemokines within individual cells that produced IFN-γ and/or TNF-α, including the cell repair cytokine amphiregulin, the chemokine MIP-1α, and the Th2 cytokines IL-4 and IL-13. Application and adaptation of this CyTOF panel to study a variety of disease states, including cancer, autoimmunity, and other chronic conditions, as well as to profile cell therapy products, will help reveal novel disease pathways and therapeutic targets.

### Panel Development

#### Overall Strategy

We developed this 50-marker mass cytometry panel to broadly investigate phenotypic shifts and functional responses of canonical peripheral blood mononuclear cell (PBMC) populations, including lymphocytes (T cells, B cells, and NK cells), myeloid-derived cells, and their distinct subsets (γδ T cells, regulatory T (Treg) cells, iNKT cells, MAIT cells, etc.). We intend to provide an optimized and validated panel as a tool for researchers to characterize the functional diversity of human immune cells across disease states and the therapeutic landscape. To comprehensively evaluate this panel, we approached optimization using a stepwise assessment strategy. This involved carefully assigning metal-antibody pairings based on (1) relevant biological abundance of each marker, (2) sensitivity of each metal, and (3) reducing potential spillover. Next, we titrated and evaluated each marker’s performance in the panel. When issues were identified (i.e., signal overlap into other channels), we adjusted and re-evaluated the panel accordingly to achieve optimization.

#### Marker Selection

The backbone of this panel includes canonical markers for identifying major human immune cell lineages. We included standard markers to identify T cells (CD3, CD4, CD8), B cells (CD19, IgD), NK cells (CD56), and myeloid-derived cells (CD14, CD11c). We were mostly interested in the functional subsets of populations within the T cell compartment and therefore selected markers to identify well-characterized phenotypic and descriptive surface markers, as well as functional cytokines and chemokines. Unconventional T cells required additional lineage markers to identify γδ T cells (TCR Vγ9, TCR Vδ1, TCR Vδ2), iNKT cells (TCR Vβ11, TCR Vα24), and MAIT cells (TCR Vα7.2, CD161). To characterize antigen exposure, we included phenotypic markers to identify naive T cells and memory T cells (CD45RA, CD27, CD45RO). We also included several phenotypic markers to assess T cell activation (CD25, CD28, PD-1), and additional markers to further subset and assess the state of the major cell lineages (CCR6, CD27, CD38, CD127, CD57, TIGIT). The remaining 24 markers included in our panel were intracellular targets to evaluate the functional state of PBMCs. We included intracellular markers to measure Th1 (IL-2, IFN-γ, TNF-α), Th2 (IL-4, IL-5, IL-13, Amphiregulin),^4–6^ Th17/22 (IL-17A, IL-17F, IL-21, IL-22), Th9 (IL-9), and regulatory responses (IL-10, TGF-β, IL-35). We included CD107a to show the degranulation capacity of immune cells in response to stimuli, as well as functional markers to assess myeloid regulation (GM-CSF, IL-3).^7^ To measure innate immune responses, we included pro-inflammatory cytokines and chemokines (IL-1β, IL-8, MIP-1α, MIP-1β).^8^ Finally, we included markers to directly measure cytotoxic activity (perforin, granzyme B).^9^

#### Metal Pairings

Antibody-metal pairings were determined by a number of factors, including but not limited to (1) relevant biological expression levels under both unstimulated and stimulated conditions, (2) sensitivity of the metals, (3) isotopic impurities, and (4) spillover signal from oxides, metal impurity, and adjacent channels.

#### Antibody Conjugations

Custom metal-tagged antibodies were conjugated using the Maxpar™ X8 Multi-Metal Labeling Kit (Standard BioTools) following the manufacturer’s instructions or through Custom Conjugation Services (Standard BioTools). Briefly, metal-chelating polymers were loaded with the metal solution and purified. Carrier-free purified antibodies were partially reduced and added to the polymer-metal complex for conjugation. Each metal-conjugated antibody was quantified and resuspended to a final concentration of 0.5 mg/mL. Please see **Supplemental Table 1** for detailed panel information.

#### Panel Assessments

All markers were evaluated using human PBMCs from healthy donors in both unstimulated and stimulated conditions to (1) accurately measure inducible marker expression and (2) ensure optimal concentrations selected for lineage markers performed across conditions. To determine the optimal concentration for each antibody, all markers were thoroughly titrated. The first phase involved titrating the major lineage markers (CD14, CD19, CD11c, CD56, CD3, CD4, CD8) to determine the optimal concentration required to achieve an appropriate signal range within each canonical population under both unstimulated and stimulated conditions (**Figure S3**). This is completed using a 5-tube titration method, in which each antibody is stained (surface or intracellularly), at 5 serially diluted concentrations, and a dilution curve is created showing signal intensity across dilutions. The dilution that demonstrated the best signal resolution with minimal background or spillover signal was selected as optimal. The second phase of the titration plan involved following the same 5-point dilution curve with the remaining 43 markers, which were titrated simultaneously as a single multiplexed cocktail using the major lineage markers listed above as counterstains (**Figure S3-4**). This approach allowed us to reliably gate on canonical populations prior to evaluating phenotypic and functional markers, and to comprehensively assess signal overlap across the full panel. Although signal overlap is far less of a concern than in fluorescence flow cytometry, it is still a necessary consideration in panel development. Assessing signal overlap focused on factors affecting the primary mass channel (M), including: (1) M+1 and M-1, neighboring channels, for isotopic impurities of the metal and abundance sensitivity of the instrument, and (2) M+16, corresponding to potential oxide species formations, mostly relevant for lanthanides, which appear as signal 16 mass units heavier than the primary isotope. In addition to the neighboring channels and oxide formations, we also evaluated potential isotopic impurities by examining the cross-section of the antibody of interest and all other antibodies paired with different isotopes of the same element. Please see **Supplemental Table 1** for information on titration details and optimal selections.

#### Stimulation Optimization

Our panel included an unprecedented number of intracellular targets that required potent stimulation to elicit a clear and measurable signal. Therefore, we used a combination of common stimuli to induce a broad and diverse cellular response. Frozen human PBMCs from multiple healthy donors were thawed, rested for a minimum of two hours at 37°C, and cultured with phytohemagglutinin (PHA) and lipopolysaccharide (LPS) overnight (∼16-18 hours), followed by additional stimulation with Phorbol 12-myristate 13-acetate (PMA) and ionomycin (I) in the presence of brefeldin A and monensin for 3-4 additional hours. Unstimulated cells from the same donors were rested at 37°C throughout the stimulation period. PHA and LPS were used to promote sustained activation of lymphoid and myeloid populations, respectively, while PMA and ionomycin provided a strong, receptor-independent stimulation of T cells and other cell types to maximize intracellular signaling and cytokine accumulation. Following stimulation, cells were washed, counted, and stained with surface and cytoplasmic antibodies as described in the Supplemental Information. Samples were acquired on a CyTOF XT system and normalized with EQ™ Six Element Calibration Beads. Please see **Supplemental Figure 2** for a representative comparison of our stimulation to PMAi and LPS alone.

#### Panel Optimization

As described earlier, the panel assessment included evaluating the spillover signal and determining the optimal concentration for each antibody. Antibody staining was performed according to the order outlined in **Supplemental Table 1**, which details the staining sequence for all antibodies in the panel. The following antibodies showed minimal signal overlap, produced a clear, distinct positive signal, and in the case of intracellular markers, a stimulation-specific signal: CD57 (89Y), CD107a (106Cd), IL-3 (110Cd), IFN-γ (116Cd), CCR6 (139La), CD8 (142Nd), CD19 (145Nd), CD14 (146Nd), IL-5 (147Sm), CD56 (149Sm), TCR Vδ2 (141Pr), CD45RA (143Nd), IL-22 (150Nd), CD161 (151Eu), CD25 (153Eu), CD27 (155Gd), PD-1 (156Gd), CD11c (158Gd), GM-CSF (159Tb), CD28 (160Dy), IL-13 (169Tm), TCR Vδ1 (174Yb), TCR Vα24 (176Yb), Granzyme B (198Pt).

Upon stimulation, certain surface receptors are known to internalize or be downregulated. To preserve detection of key lineage markers under these conditions, CD3 (170Er) and CD4 (144Nd) were stained during the intracellular staining step.

The following antibodies showed a spillover signal into either the M+1 or M-1 channel within the log-fold detection range, which was resolved by using a more dilute antibody concentration of either the antibody in question or the comparison antibody: IL-9 (111Cd), IL-2 (112Cd), MIP-1α (113Cd), Vγ9 (115In), IL-35 (154Sm), CD38 (161Dy), Amphiregulin (162Dy), CD45RO (163Dy), IL-10 (165Ho), IL-4 (171Yb), IL-21 (172Yb), TCR Vα7.2 (194Pt), IgD (196Pt), IL-17A (195Pt), Perforin (173Yb), and TGF-β (164Dy).

In its functionally mature form, TGF-β is a dimeric ligand that binds to type II receptor dimers (TGF-βRII), recruits type I receptor dimers (TGF-βRI), and forms a tetrameric receptor complex at the plasma membrane; therefore, TGF-β was stained on the cell surface to measure this active protein complex.^10^

Three antibodies also showed signal overlap within the log-fold detection range due to isotopic impurities of 166Er, 167Er, and 168Er; CD127 (168Er) was observed to downregulate upon stimulation; therefore, we decided to select a more concentrated antibody titration. We also observed isotopic impurities with CD127 (168Er) when compared with IL-17F (166Er) and TIGIT (167Er), which were resolved at more dilute concentrations of the comparing antibodies.

TCR Vβ11 (148Nd) exhibited spillover into the M+16 channel within the log-fold detection range, which was resolved by using a more dilute antibody concentration.

MIP-1β (152Sm) and TNF-α (114Cd) exhibited potential isotopic impurity issues within the log-fold detection range, which were resolved by using a more dilute antibody concentration.

During the panel optimization stages, we had to reassign two antibodies to different metal channels. IL-8 (175Lu) was originally assigned to 139La, where we observed substantial oxide formation into the 155Gd channel. After attempts to resolve this issue by increasing the dilution used, we ultimately decided to reassign the antibody marker to a new metal, 175Lu. After re-metaling this marker, we observed a minor signal overlap, less than a log-fold, into the M+1 channel, which was resolved to an acceptable level with a more dilute antibody concentration. IL-1β (209Bi) was originally assigned to 173Yb, resulting in spillover into adjacent channels upon stimulation. We reassigned this antibody marker to 209Bi, where it showed no spillover and produced a clear, distinct, stimulation-dependent positive signal.

### Data Analysis

Events were first enriched for live immune cells using the gating strategy shown in **Supplemental Figure 1**. A representative analysis of the complete 50-parameter panel from live lymphocytes is shown (**Figure 1**). First, we identified CD19+ and/or IgD+ from live intact singlets to isolate the B cell lineage population (**Figure 1A**). Moving forward, we used the CD19-IgD-subset; we gated on the myeloid-derived immune lineage by identifying CD14+ and/or CD11c+ cells. From CD14-CD11c-cells, we gated on CD3+ cells, including CD56+/-cells, identifying T cells, and CD3-CD56+/-cells, identifying NK cells and Innate Lymphoid Cells (ILCs). From the identified T cell population, we next gated on Vγ9+ and/or Vδ1+ or Vδ2+ cells to identify γδ T cells. Once we identified our γδ T cells, we used the non-γδ T cell population to identify further subsets of canonical and non-canonical T cells (CD4+, CD8+, iNKTs, and MAITs). Next, we gated on the remaining surface markers, most of which also include additional phenotypic markers. Cytokine detection is shown in representative bivariate plots compared to the corresponding unstimulated control, with gating on stimulated CD3+ CD56+/-T cells (**Figure 1B**), cytotoxic CD8+ T cells (**Figure 1C**), and monocytes and dendritic cells (**Figure 1D**).

**Figure 1.**
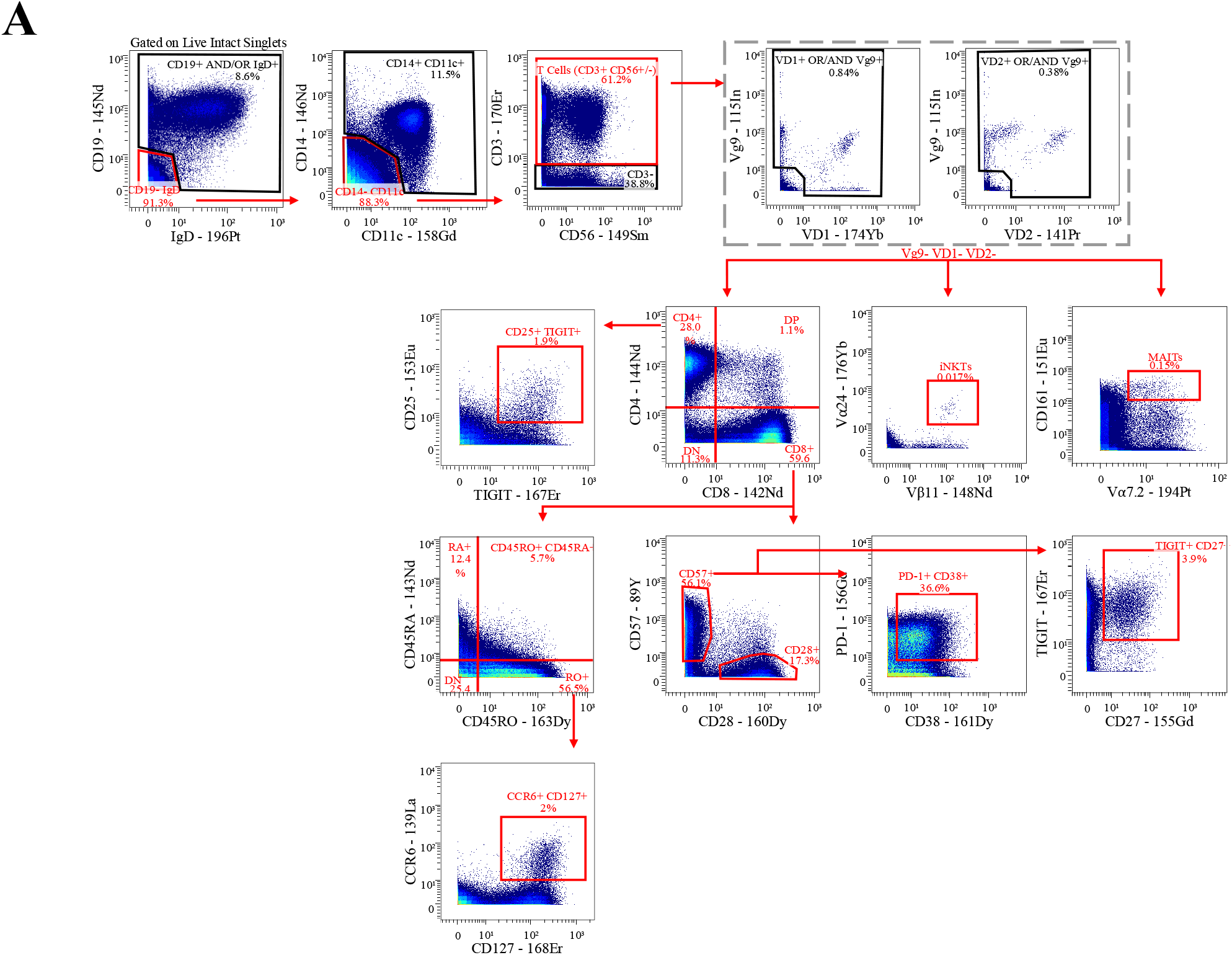
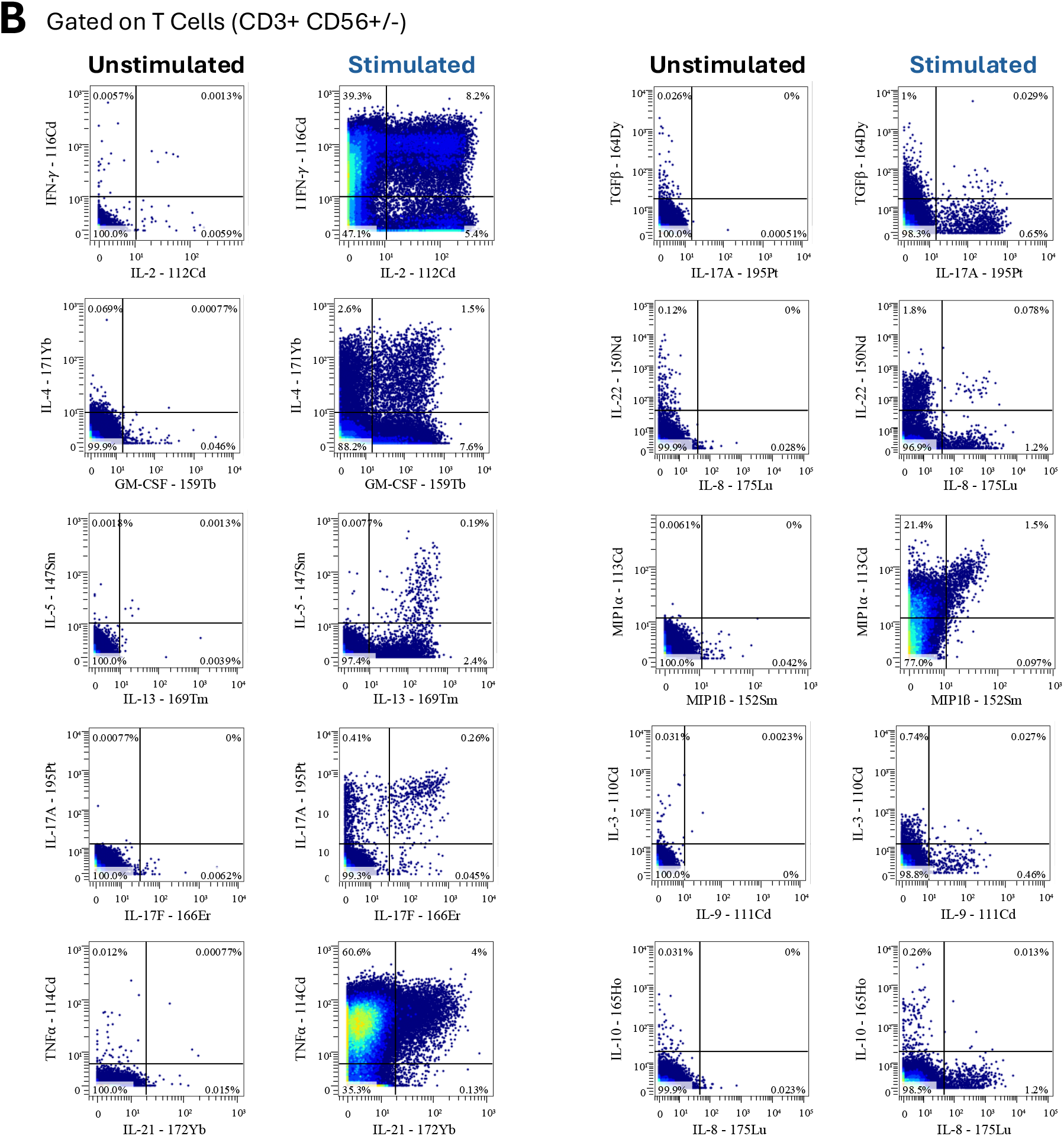
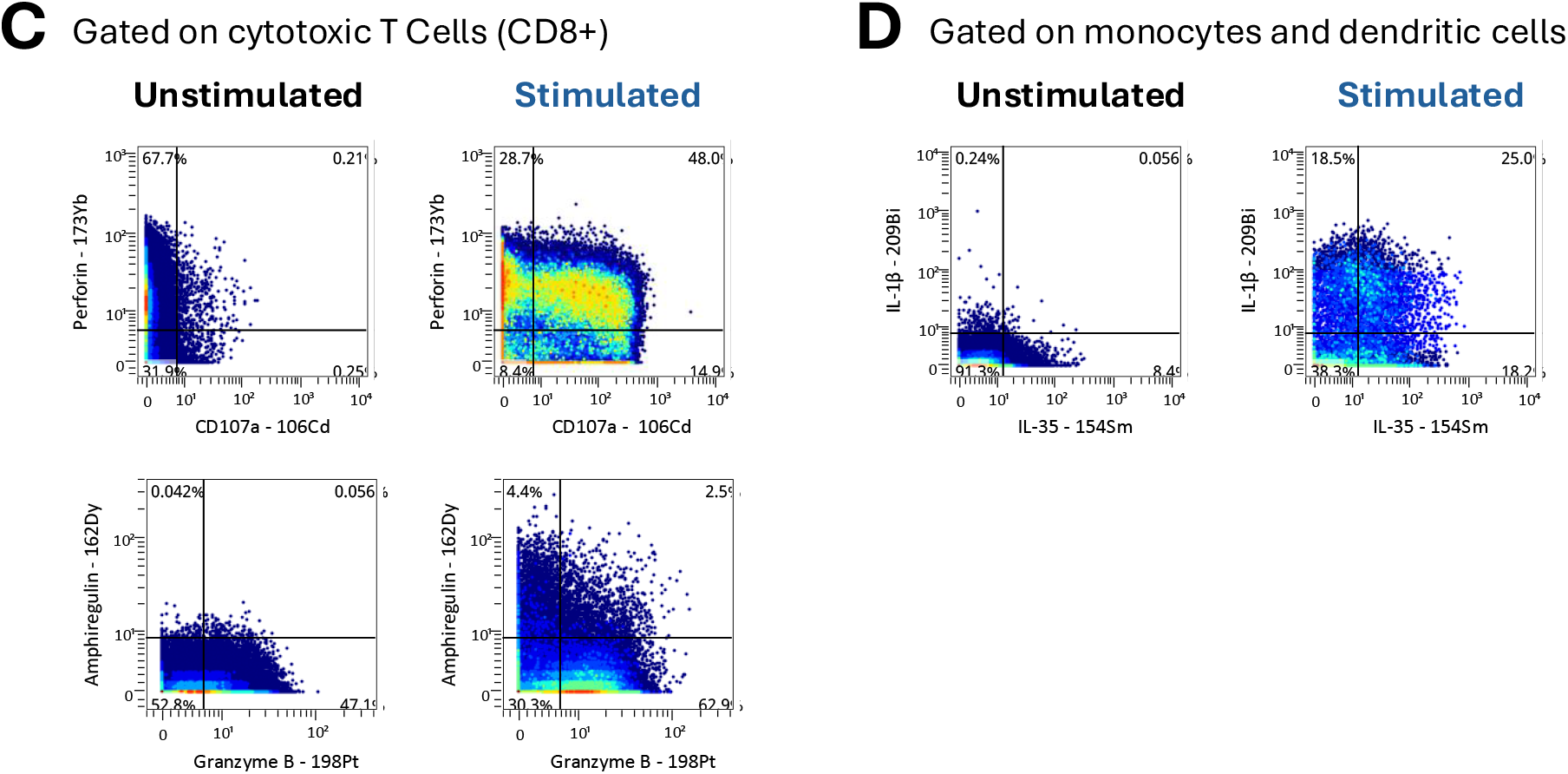
Manual gating strategy. Healthy PBMCs were utilized to showcase the panel. First, clean-up gating was successfully applied to identify live intact single cells (see Supplemental Figure 1). Arrows indicate parent to child plots in the direction of the arrow. (A) CD19+ and/or IgD+ were gated from live intact singlets to isolate the B cell lineage population. Of the CD19-IgD-subset, we identified myeloid-derived cells using CD14+ and/or CD11c+ events. From CD14-CD11c-cells, we gated on CD3+ cells, including CD56+/-cells, identifying T cells. From the identified T cell population, we next gated on Vg9+ and/or Vδ1+ or Vδ2+ cells to identify γδ T cells. The non-γδ T cell population was used to identify unconventional T cells, iNKTs using Vα24 and Vβ11, and MAITs using Vα7.2 and CD161. We also used the non-γδ T cell parent gate to identify conventional T cell lineages. We used a quadrant gate to identify CD4+, CD8+, double-negative, and double-positive populations. We used the CD4+ quadrant to identify cells expressing both CD25 and TIGIT. From there, we used the CD8+ quadrant of the parent plot to distinguish CD45RO from CD45RA, thereby showcasing memory vs naïve cells. From the CD45RO+ quadrant, we gated on CCR6 and CD127-positive events. The CD8+ parent quadrant was also used to distinguish between CD57 and CD28 co-expressing cells. From there, we used the CD57 gated population to identify PD-1 and CD38 co-expressing cells as well as TIGIT and CD27 co-expressing cells. Functional readouts are shown across canonical immune linages, including **(B)** T cells (CD3+ CD56+/-) **(C)** T cells (CD8+), and **(D)** Monocytes and dendritic cells.

To determine how well this panel can identify and define memory T cells across varying functional lineages, we projected multiparameter data from live CD45RO+ T cells (enriched for antigen-experienced T cells) into a 2D space using opt-SNE.^13^ We then clustered the same multiparameter dataset using Phenograph^11^, and identified 23 clusters with distinct surface phenotypes and intracellular readouts (**Figure 2A**). Well-established functional lineages were confirmed, including Th17, Th2, and Tc1 cells (clusters 20, 14, and 22, respectively), providing important validation of the panel’s accuracy (**Figure 2B**). As expected, not every intracellular target was detected in memory T cells; however, monocytes (defined as CD14^var^ CD11c+) produced IL-1β, IL-35, and IL-8 (**Figure 2B**), as well as MIP-1α and MIP-1β. In summary, this analysis visualized the vast, robust, and complex responses captured by our comprehensive functional panel.

**Figure 2.**
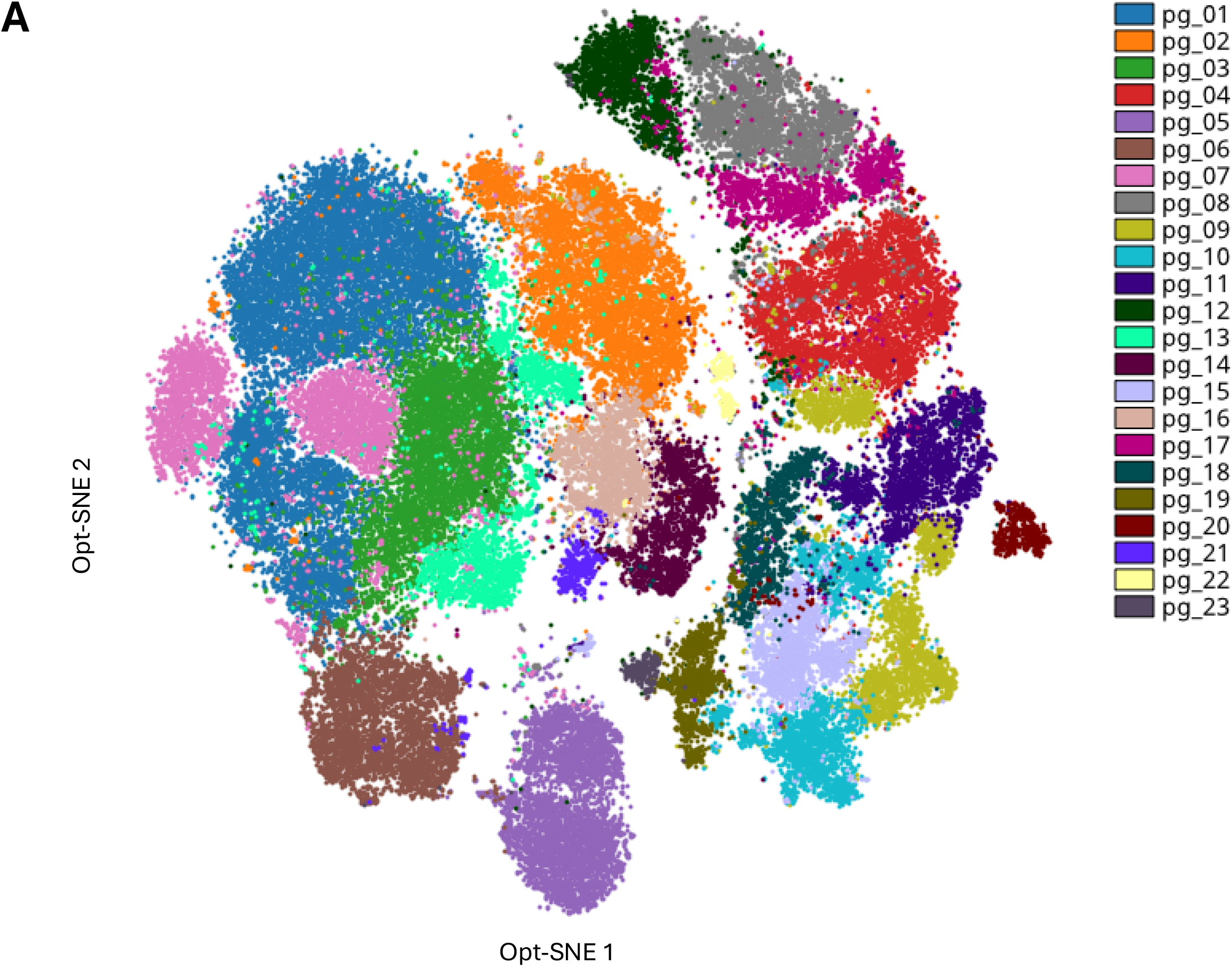
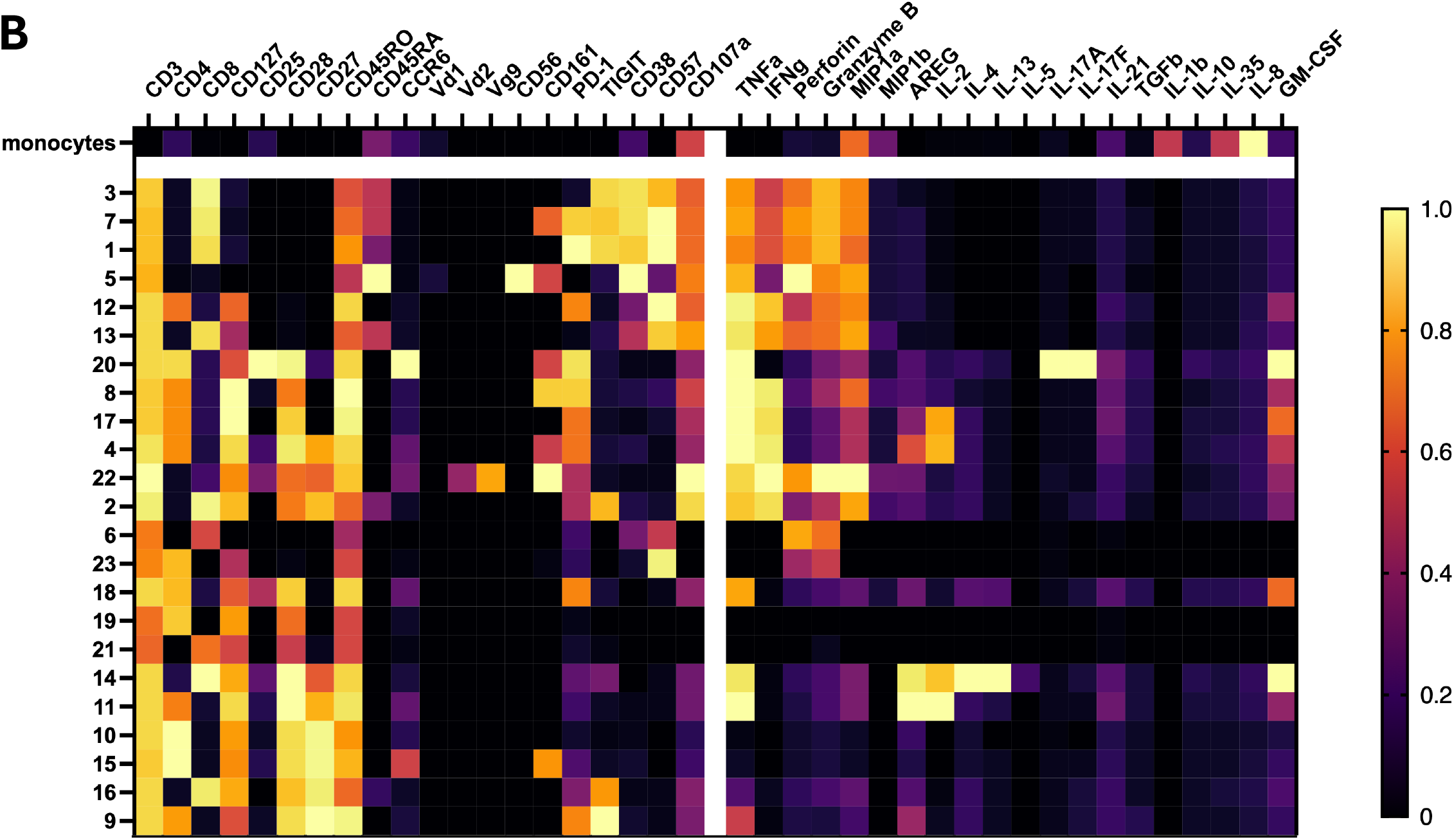
High-dimensional single-cell data visualization. (**A**) Live CD45RO+ T cells were used to generate an opt-sne visualization map. Using the same population, we clustered the dataset using Phenograph, identifying 23 distinct clusters (pg_01-pg_23), (**B**) The same phenograph clusters were visualized as a heatmap for additional comparative analysis alongside monocytes.

Next, we tested the utility of this panel for profiling T cell function in response to physiological stimulation; to do this, we used a CEF peptide pool, consisting of a total of 32 peptides from HLA class I-restricted T cell epitopes from the three viruses. Healthy PBMCs were cultured overnight with or without these peptides, in the presence of protein transport inhibitors, and cells from both culture conditions were stained with a reduced version of the panel, analyzed on a CyTOF Helios instrument, and total cell populations were visualized using opt-SNE. A clearly distinct population of cells producing TNF-α, IFN-γ, or IL-2 was identified, comprising ∼7.7% of T cells, within the normal range of CEF responses reported by others.^12^ Also, the vast majority of CEF-peptide-specific T cells produced both TNF-α and IFN-γ, consistent with a previous report.^12^ Many of these Th1 cytokine+ T cells also produced the chemokine MIP-1α, and, to our surprise, many also produced the cell repair and TGF-β-inducing cytokine amphiregulin (**Figure 3B**).^4^ In addition, a small but clearly detectable subset of these cells produced the Th2 cytokines IL-4 and IL-13, revealing individual virus-specific ‘Th0’ cells that simultaneously produce both Th1 and Th2 cytokines (**Figure 3B**).^13–15^

**Figure 3.**
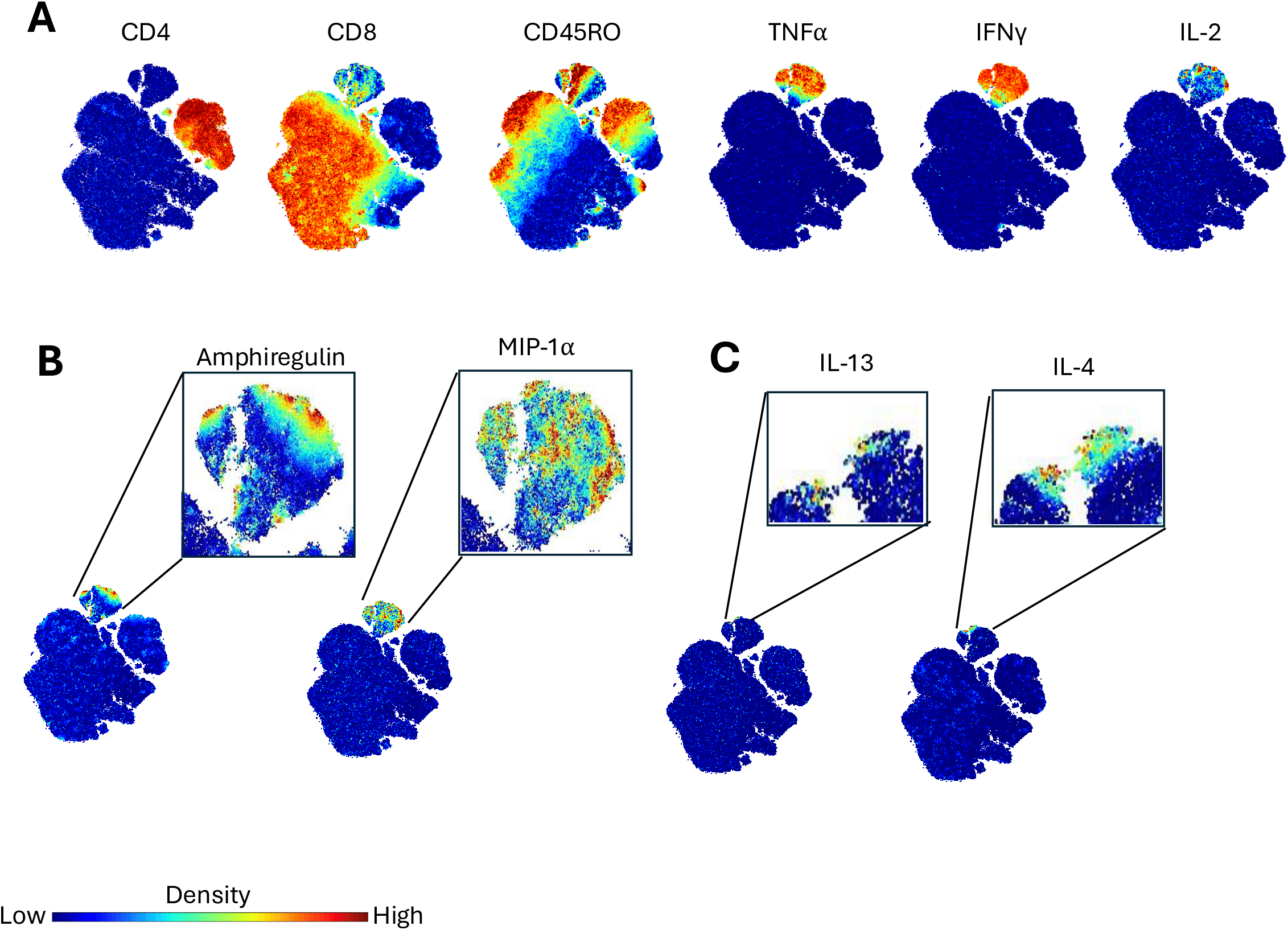
Opt-sne maps of CEF-stimulated healthy PBMCs. Density is visualized from low to high, with cool to warmer-colored points on each plot, respectively. (**A**) Canonical CD4, CD8, and CD45RO are shown along with Th1-associated cytokines TNF⍰, IFNγ, and IL-2. (**B**) The Th1 cytokine-producing cells also produce amphiregulin and chemokine MIP-1 ⍰. (**C**) Within this population, there is a rarer subset of cells also producing Th2-associated cytokines IL-13 and IL-4.

## Discussion

We have developed a 50-marker human mass cytometry panel that enables the detection of many populations of immune cells (including both unconventional and adaptive T cell populations, B cells, NK cells, and monocytes) that produce/express combinations of 24 cytokines, chemokines, and other functional readouts. Specifically, this panel allows simultaneous detection of both cytokines and effector molecules commonly included in fluorescent cytometry panels, such as TNF-α, IFN-γ, IL-17A, IL-2, IL-4, and Granzyme B, and also many more functional readouts, including IL-1β, IL-8, IL-5, IL-10, IL-13, IL-21, amphiregulin, MIP-1α, and MIP-1β (**Figures 1-3**). This work and our previous study^3^ both support the ability of mass cytometry to enable highly multiplexed intracellular detection at a scale that remains difficult for fluorescence-based platforms, particularly for exploratory cytokine and effector-molecule analysis. In both the present study and our previous work, every intracellular cytokine or effector molecule confirmed by an orthogonal method was also detectable by mass cytometry. The combination of broad multiplexing and reduced dependence on predicted marker co-expression, especially when paired with computational single-cell analysis methods, makes this platform well-suited to unbiased detection across diverse functional states of human immune and other cells.

Mass cytometry does not have limitations found in fluorescence-based flow cytometry that can stranglehold panel size and resolution, as discrete reporters are resolved by atomic mass with minimal channel overlap, and autofluorescence-derived background is nonexistent. Thereby, this panel enables exploratory investigation of a broad set of cytokines and effector molecules without the need to impose prior assumptions about co-expression patterns on the panel design. The relevant cytokine combinations are not known in advance; therefore, unexpected co-expression is a target of discovery rather than a design problem to be avoided. This is a particularly important approach when investigating functional states that are incompletely characterized, including the selective chemotaxis of different cell populations, active abrogation of inflammation, as well as cell debris removal and cell repair programs to return microenvironments to homeostasis after immunogens are cleared/pathogens are contained. Large fluorescence panels remain highly dependent on anticipated marker expression patterns: fluorochrome choices are interpretable only when expected co-expression and spectral overlap are considered together. Unexpected co-expression of markers assigned to overlapping fluorochromes can therefore become a major source of ambiguity, particularly in exploratory panels where the relevant expression patterns are not known a priori. This problem is especially pronounced for intracellular cytokine detection, where stimulation and fixation increase autofluorescence, fixation can degrade fluorescent labels, and cytokine signals are often dim. Therefore, mass cytometry was the optimal platform to create a panel for unbiased, highly multiplexed proteomic functional analysis of individual human immune cells.

The overwhelming use of fluorescent intracellular cytokine panels to assess human T cell function has predominated human immunology for the last three decades; this has resulted in dim or technically challenging readouts to sometimes be underreported and/or misinterpreted as absent; this has stunted the field. For example, we recently reported reproducible detection of IL-13-producing peripheral blood T cells after mitogen stimulation,^3^ whereas some prior fluorescence-based studies reported little or no IL-13-producing cells under comparable conditions.^19^ These discrepancies underscore the vital importance of routine use of both positive controls within experimental runs to confirm assay detection capability, and unfortunately, researchers must possess an ardent skepticism of the performance claimed by the manufacturers of both cytometry reagents and analysis instruments.

A further advantage of mass cytometry includes the relative flexibility of antibody-metal conjugation. CyTOF antibody reagents can be generated by end users using standardized conjugation kits and compatible purified antibodies, allowing exploratory panel expansion with substantially fewer constraints from spectral overlap, anticipated marker co-expression, and the limited portfolios of preconjugated fluorescent antibodies available from manufacturers. By contrast, in-house fluorochrome conjugation options are relatively limited, and custom conjugation to newer dyes often requires bulk ordering of custom reagents, with no guarantee that a given fluorochrome–marker combination will be compatible with the rest of the panel. The panel is modular, and many antibody-metal conjugates can be generated in-house; it can be adapted to emerging biological questions and to investigate more nuanced or previously understudied phenotypes.

Future adaptations to this panel could include markers associated with specialized pro-resolving mediator pathways,^16^ for example, and a more immediate application would be to use this platform to improve the functional assessment of cell therapies. Our preliminary data indicate substantial functional diversity in response to CAR antigen stimulation (data not shown), and we predict that the limited efficacy of some Treg cell therapies may be explained, at least in part, by a scarcity of genuinely immunosuppressive T cells within individual products. Our platform can be used to assess whether suppressive functional states are sufficiently represented within Treg therapy products and whether ‘saboteurs’ (i.e., pro-inflammatory T cells) are subverting the efficacy of the drug.

It is important to note that many intracellular readouts can be resolved comparably by mass cytometry and fluorescence cytometry.^3^ Indeed, our group has detected IL-5, IL-10, and IL-13 in small fluorescence-based panels when these cytokines were assigned to bright small organic-molecule fluorochromes like Alexa Fluor 488 and Alexa Fluor 647, and when the panel was redesigned to bring the spectral interference to an absolute minimum (data not shown). The limitation, as we can discern from our results to date, is therefore not detection *per se*, but scale. The number of such favorable fluorochrome channels is extremely limited, and panels built around these constraints cannot easily accommodate deep phenotyping or broad functional profiling. Thus, while the 50-marker CyTOF panel is well-suited for exploratory discovery and broad co-expression analysis, smaller fluorescence panels remain valuable for targeted intracellular functional assays designed to follow up and focus on specific pathways or candidate readouts.

Some functional phenotypes we detected in healthy PBMCs were rare even after potent mitogenic stimulation (for instance, IL-5, IL-13, and IL-9). However, low frequency in this reference setting does not indicate poor assay performance or limited utility. These readouts showed robust stimulation-specific induction, supporting their detectability and biological interpretability within the panel. Their rarity in healthy PBMCs may instead reflect the composition and differentiation history of the sampled population. In disease settings, tissue isolates, antigen-specific enriched populations, or cell therapy products, the same readouts may identify substantially expanded functional subsets and therefore provide information that would be missed by panels restricted to more abundant cytokine responses.

This platform could therefore provide a practical framework for assessing functional heterogeneity within cell therapy products and for comparing products across batches or donors. Beyond cell therapy, broad single-cell cytokine profiling may help resolve questions of plasticity and lineage commitment in antigen-specific memory cells, innate-like T cells, and other immune populations. Future panel iterations can also incorporate newly identified effector molecules emerging from proteomic or transcriptomic discovery.

Together, these results establish a high-dimensional 50-parameter panel for single-cell comprehensive profiling of human immune cell function and surface phenotype. By combining broad immune subset identification with simultaneous detection of 24 cytokines, chemokines, and effector molecules, this panel enables exploratory analysis of functional co-expression patterns that are difficult to capture using narrower or more hypothesis-constrained approaches. Our panel is particularly relevant for studying chronic inflammation, vaccine responses, and therapeutic interventions, including CAR-T and other cell therapy products.

### Similarity to Published OMIPs

To our knowledge, **OMIP-060**,^17^ which focuses on identifying immune cell compartments, is currently the most similar to our panel in purpose. However, our panel, which utilizes mass cytometry rather than spectral-based flow cytometry, expands the phenotypic capacity to include MAIT cells and additional innate populations, as well as additional functional molecular targets spanning various immunological processes and pathways. **OMIP-034**^18^ is a similarly designed panel for comprehensive profiling of immune cells using mass cytometry in response to immunomodulatory drugs, but it has limitations due to the absence of functional markers. To date, no other OMIP is designed to investigate both the phenotypic and functional capabilities of the peripheral immune system to this breadth and diversity.

## Author Contributions

**Laura C. Polanco:** investigation, methodology, writing – original draft, writing – review and editing, formal analysis. **Michael J. Cohen:** investigation, methodology, writing – original draft, writing – review and editing. **Lauren Tracey:** investigation, methodology, writing – original draft, writing – review and editing. **Christina Loh:** conceptualization, investigation, writing – review and editing, supervision. **Erika L. Smith-Mahoney**: experimentation, methodology, review, and editing. **Amedeo J. Cappione:** conceptualization, investigation, writing – original draft, methodology, writing – review and editing. **David King:** conceptualization, investigation, writing – original draft, methodology, writing – review and editing. **Anna C. Belkina:** conceptualization, investigation, formal analysis, supervision, writing – original draft, writing – review and editing. **Jennifer E. Snyder-Cappione:** conceptualization, investigation, supervision, formal analysis, writing-original draft, writing – review and editing.

## Notes

### Competing Interest Statement

The authors have declared no competing interest.

### Funding

- NIH/National Institute on Aging: 5R01AG065050-05

