## Supplemental Figures for "A 50-marker mass cytometry panel to expand analysis of the functional breadth of human immune cells"

**STAINING PROTOCOL**

*Materials*

- Greiner Bio-One CELLSTAR Cell-Repellent Surface Cell Culture 24-Well Plates (Greiner Bio-One, 07-000-683)
- Maxpar PBS (Standard BioTools, 201058)
- Maxpar Cell Staining Buffer (Standard BioTools, 201068)
- Maxpar Fix and Perm Buffer (Standard BioTools, 201067)
- Maxpar Perm-S Buffer (Standard BioTools, 201066)
- Gibco™ RPMI 1640 Medium, HEPES (Gibco, 22400089)
- Corning® Fetal Bovine Serum, 500 mL, Regular, USDA Safety Tested (Heat Inactivated) (Corning, 35-011-CV)
- Gibco™ Penicillin-Streptomycin (10,000 U/mL) (Gibco, 15140122)
- Cell Activation Cocktail without Brefeldin A (500X) (BioLegend, 423301)
- Brefeldin A (1000X) (Sigma, B7651)
- Monensin (Sigma, M5273)
- Invitrogen™ eBioscience™ Lipopolysaccharide (LPS) Solution (500X) (Invitrogen™, 00497693)
- Phytohemagglutinin PHA-M, lyophilized powder (Sigma, L2646-10MG)
- Human TruStain FcX^TM^ (BioLegend, 422301)
- Pierce™ 16% Formaldehyde (w/v), Methanol-free, 10 x 10 mL (Thermo Fisher, 28908)
- Cryopreserved PBMCs were obtained from whole blood isolation and frozen in aliquots or purchased (Stem Cell Technology, 70025.1)
- Cell-ID^TM^ Intercalator-IR (Standard BioTools, 201192C)
- Heparin in PBS – 10KU/mL (Sigma, H3149-100KU)

*Buffer Preparation:*

- R10 Media: Prepare R10 cell culture media using sterile Gibco™ RPMI 1640 Medium with 10% FBS and 1% Penicillin-Streptomycin.
- Prepare the surface staining antibody cocktails (pre-stain 1, pre-stain 2, and bulk) and the intracellular antibody staining cocktail fresh on the day of their use with a final volume of 100 µL per sample. CD107a will be added during the stimulation protocol.
- 1.6% FA: Prepare a fresh 1.6% Formaldehyde solution from the 16% stock ampule, enough for 1 mL per sample. Dilute 1 part of the filtered stock with 9 parts of the Maxpar PBS.
- Intercalation solution: Prepare 1 mL of cell intercalation solution for each unique sample by adding Cell-ID Intercalator (191/193 Iridium) into Maxpar Fix and Perm Buffer (1:500 dilution of the 12.5 µM stock solution for a 25 nM final concentration) and vortex to mix. Prepare fresh for same-day use.
- Heparin-block solution: Prepare a working stock of Heparin-blocking solution by diluting the Heparin-block stock from 10KU/mL to 100 U/mL in Maxpar Perm-S Buffer.

*Sample Thawing & Stimulation*

**Experiment Day 1:**

1. Prepare a single 50 mL conical tube with 20 mL of cold R10 media for each unique PBMC donor. Carefully thaw vial(s) in a wet or dry bath set to 37 °C for 30-60 seconds. Transfer the PBMCs to the prepared conical containing R10 media. Add up to 50 mL of cold R10 to each conical tube(s). Centrifuge the cells for 10 minutes at 300 x *g*.
2. Decant the supernatant into the waste bucket. Add 20 mL of R10 to the conical tube(s). Gently invert the conical tube(s) 3-4 times to resuspend the pellet(s).
3. Count PBMCs. Set samples on ice while counting. Adjust cells to 6e6 cells/mL using R10.
4. Seed PBMCs, 500 µL per well in a 24-well plate.
5. Allow cells to rest for at least 2 hours in a CO_2_ incubator set to 37 °C, 5% CO_2_, and 95% relative humidity.
6. Prepare working stocks of stimulation reagents, PHA (1 mg/mL stock) and LPS (2.5 mg/mL), in R10 media at dilutions of 1:10 (PHA) and 1:25 (LPS) to achieve a final *in vitro* concentration of 1 µg/mL for each. Add 5 µL of each working stock to the stimulation wells.
7. Place in CO_2_ incubator overnight (~16-18 hours).

**Experiment Day 2:**

8. Prepare a fresh 10% bleach waste bucket.
9. Prepare a working stock of stimulation reagent Cell Activation Cocktail 500X (PMA, 25 µg/mL; Ionomycin, 500 µg/mL stock) in R10 media at a 1:10 dilution. Add 5 µL of working stock Cell Activation Cocktail to each stimulation well for a final concentration of 0.5X (PMA, 0.025 µg/mL; Ionomycin, 0.5 µg/mL).
10. Add 1.25 µL of CD107a to each well, both unstimulated and stimulated.
11. Place in the CO_2_ incubator for 1-2 hours.
12. Prepare a working mixture of Brefeldin A (10 mg/mL stock) and Monensin sodium salt (50 mg/mL stock) in R10 media at a 1:10 dilution for a final *in vitro* concentration of 5 µg/mL for each. Add 3 µL of the working mixture to each well, both unstimulated and stimulated.
13. Place in CO_2_ incubator for up to a total stimulation time of 3-4 hours. Use the addition of the Cell Activation Cocktail as the starting stimulation time.

*Cell Viability & Surface Staining*

14. Harvest cells in a 15 mL conical tube and pool wells by condition and unique PBMC donor. Count cells.
15. Wash cells by adding up to 15 mL of Maxpar Cell Staining Buffer. Centrifuge for 5 minutes at 300 x *g*. Decant supernatant into the waste bucket and resuspend the pellet in the residual volume by gently vortexing.
16. Set aside 3E6 cells per sample into a clean 5 mL Eppendorf or 15 mL conical tube for staining. Top up to 3-5 mL with Maxpar Cell staining Buffer to repeat wash.
17. Carefully aspirate supernatant and resuspend each cell pellet in 50 µL of Maxpar Cell Staining Buffer. Transfer each pellet into a 96-well V-bottom plate. Add 5 µL of Human TruStain FcX^TM^ to each resuspended sample. Gently vortex to mix the sample thoroughly. Incubate the mix for 10 minutes at room temperature.
18. Without removing the Human TruStain FcX^TM^, add 5 µL of antibody cocktail “Pre-stain 1” to each sample. Gently mix by pipetting the mixture. Incubate for 5 minutes at room temperature.
19. Add 5 µL of antibody cocktail “Pre-stain 2” to each sample. Gently mix by pipetting the mixture. Incubate for 10 minutes at room temperature.
20. Add 35 µL of the “Bulk” antibody cocktail to each sample to achieve a total staining volume of 100 µL. Gently mix by pipetting the mixture. Incubate for 30 minutes at room temperature.
21. Wash cells with 100µL of Maxpar Cell Staining Buffer. Centrifuge for 5 minutes at 300 x *g*. Gently decant the supernatant into the waste bucket. Resuspend the pellet in the residual volume by gently vortexing. Repeat wash with 200 µL of Maxpar Cell Staining Buffer.

*Intracellular Staining*

22. Add 200 µL of the prepared 1.6% FA solution to each well, then mix thoroughly by pipetting. Incubate for 10 minutes at room temperature.
23. Centrifuge for 5 minutes at 800 x *g*. Gently decant the supernatant into the waste bucket and resuspend cells in 200 µL of Maxpar Perm-S Buffer.
24. Centrifuge for 5 minutes at 800 x *g*. Gently decant the supernatant into the waste bucket. Repeat the wash and decant.
25. Add 50 µL of Heparin-blocking mix to each sample, then resuspend the cells by pipetting. Incubate for 15-20 minutes at room temperature.
26. Without removing the heparin-blocking mix, add 50 µL of the intracellular antibody cocktail to each sample. Pipette to mix. Incubate for 30 minutes at room temperature.
27. Wash cells with 100 µL of Maxpar Cell Staining Buffer. Gently vortex to mix. Centrifuge for 5 minutes at 800 x *g*. Repeat wash with 200 µL of Maxpar Cell Staining Buffer.
28. Add 200 µL of 1.6% FA solution to each sample and vortex to mix. Incubate for 10 minutes at room temperature.
29. Centrifuge cells for 5 minutes at 800 x *g*. Decant the supernatant into the waste bucket and resuspend cells in residual volume.

*Intercalation Incubation*

30. Add 200 µL of the prepared intercalation solution to each sample. Ensure all samples are well resuspended.
31. Transfer cells from the 96-well plate to 5 mL polypropylene tubes.
32. Top up each tube to 1 mL by adding 800 µL of intercalation solution.
33. Incubate the samples overnight (up to 48 hours) at 4-8 °C.

**Experiment Day 3:**

34. Thoroughly resuspend the samples by gently vortexing.
35. Centrifuge for 5 minutes at 800 x *g*. Without disturbing the cell pellet, carefully aspirate off the supernatant. Gently vortex to resuspend the cells in residual volume.
36. Wash cells with 2 mL of Maxpar Cell Staining Buffer (CSB) and gently vortex to mix. Centrifuge for 5 minutes at 800 x *g*. Carefully aspirate off the supernatant and repeat the wash.
37. Repeat steps 34-36 for a total of 2 washes with Maxpar CSB.
38. Add 2 mL of Maxpar Cell Acquisition Solution Plus (CAS+) and gently vortex to mix.
39. Filter cells through a 35 µm cell strainer into new 5 mL polypropylene tubes.
40. Reserve 10 µL from the samples to count cells
    a. Cells should be acquired at a concentration of ~ 1x10^6^ cells/mL
41. Carefully aspirate and discard supernatant
42. Leave cells pelleted at 2-8 °C in the chilled autosampler carousel until ready to be acquired.
43. Acquire data on a CyTOF XT.

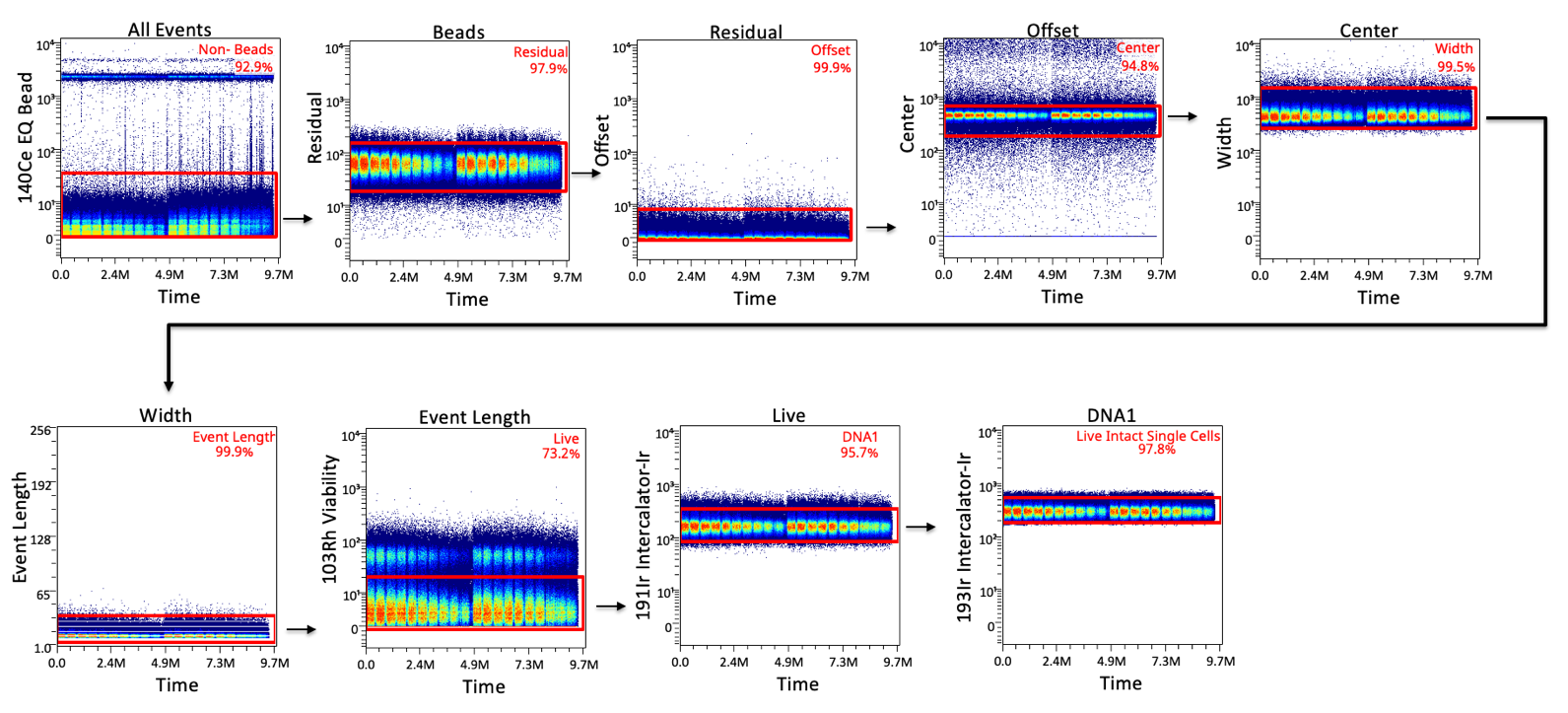
**Supplemental Figure 1. Clean up gating strategy.** Each parameter is plotted against time. Black arrows indicate the subsequent plot from the selected gate in red. Each gate is centered on the largest band. First, we used the 140Ce – “Beads” channel to gate on low intensity events to exclude calibration beads downstream. The next five gates were used to exclude debris and doublets, based on plots of Time versus residual, offset, center, width, and event length. From there, we plotted against the 103Rh-viability marker to gate on live events on the low-mid range intensity. Finally, we used DNA1 and DNA2 plots to gate on live intact single cells.

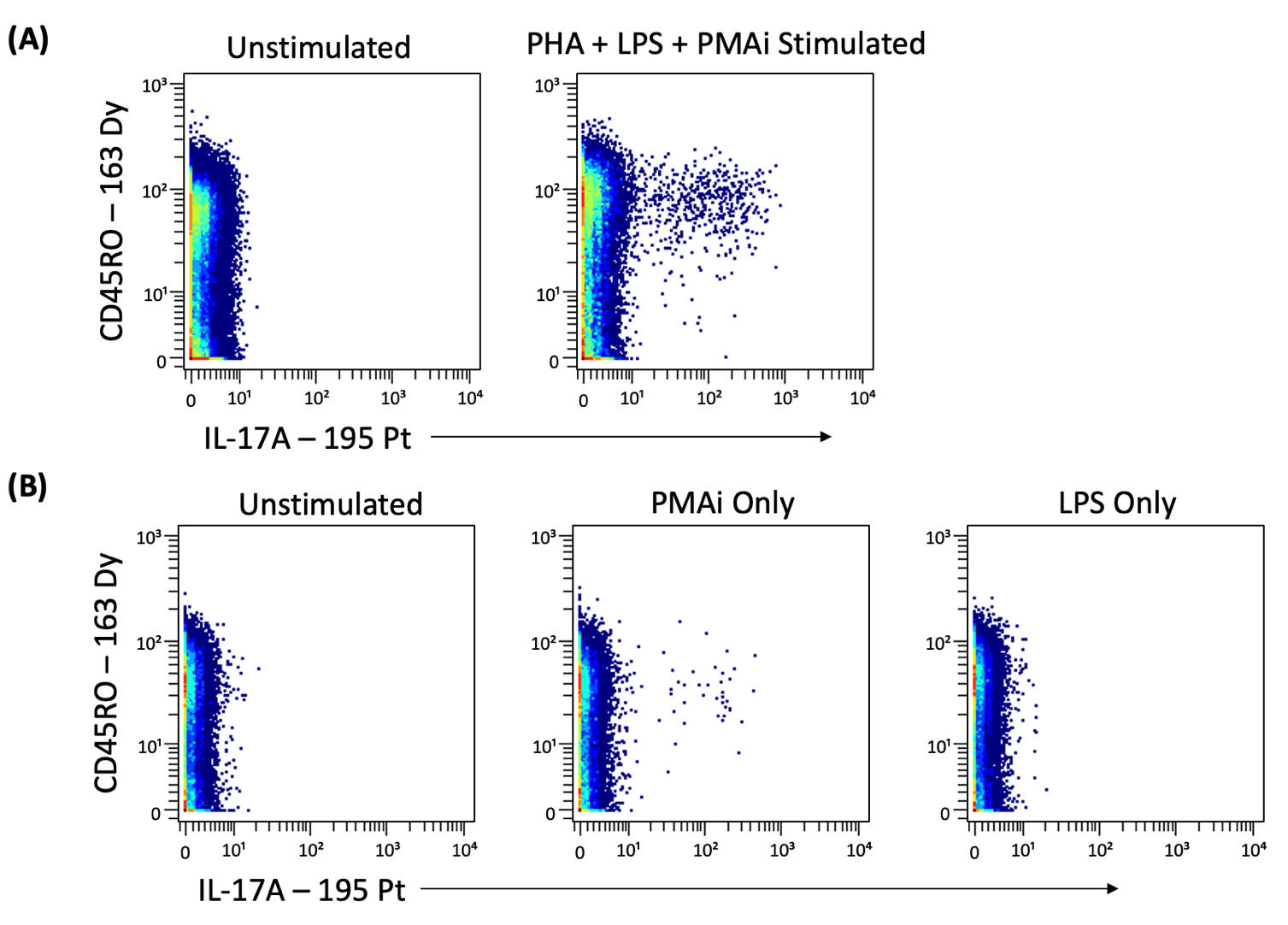

**Supplemental Figure 2. Stimulation comparison.** (A) Cells gated from CD4+ T cells demonstrating IL-17A expression upon staggered stimulation protocol compared to an unstimulated control. (B) Cells gated from CD4+ T cells demonstrating IL-17A expression upon either PMA/ionomycin or LPS overnight stimulation compared to an unstimulated control.

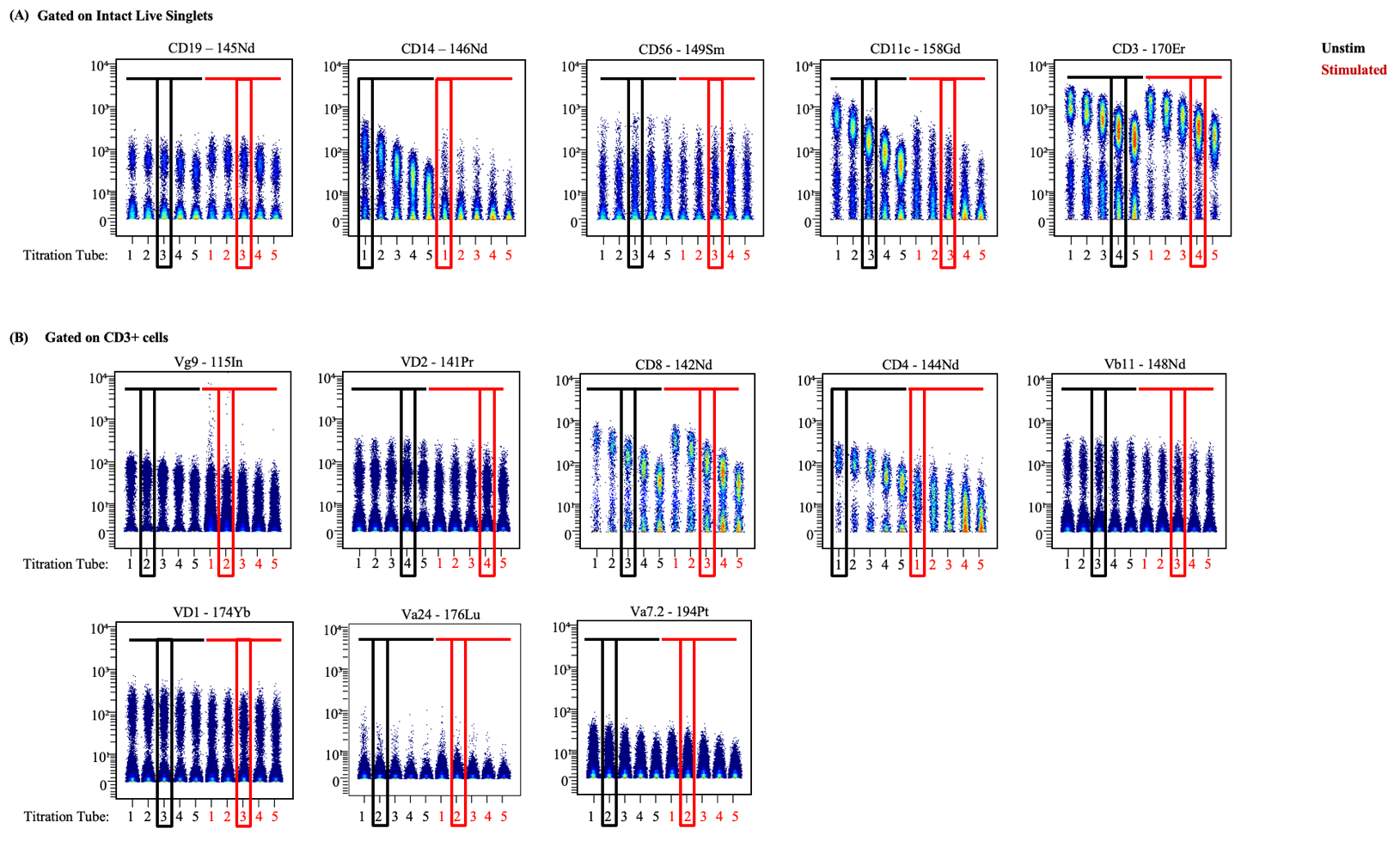

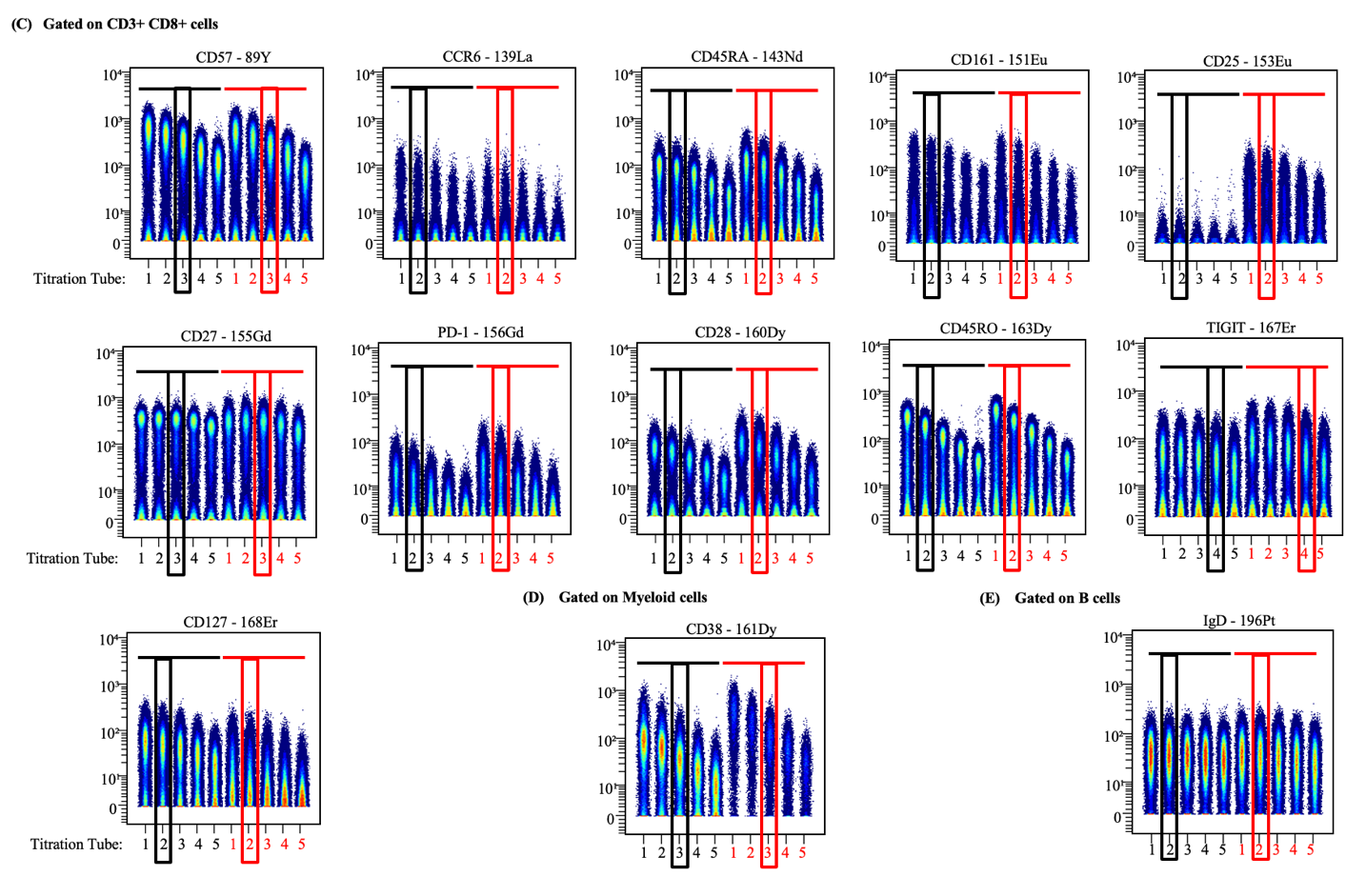
**Supplemental Figure 3. Surface marker titrations.** All antibodies were titrated on PBMCs from healthy human donors under both unstimulated and stimulated conditions. Black boxes indicate the selected optimal titration within the unstimulated samples (T1-5), and the red boxes indicate the same selection within the stimulated samples (T6-10).

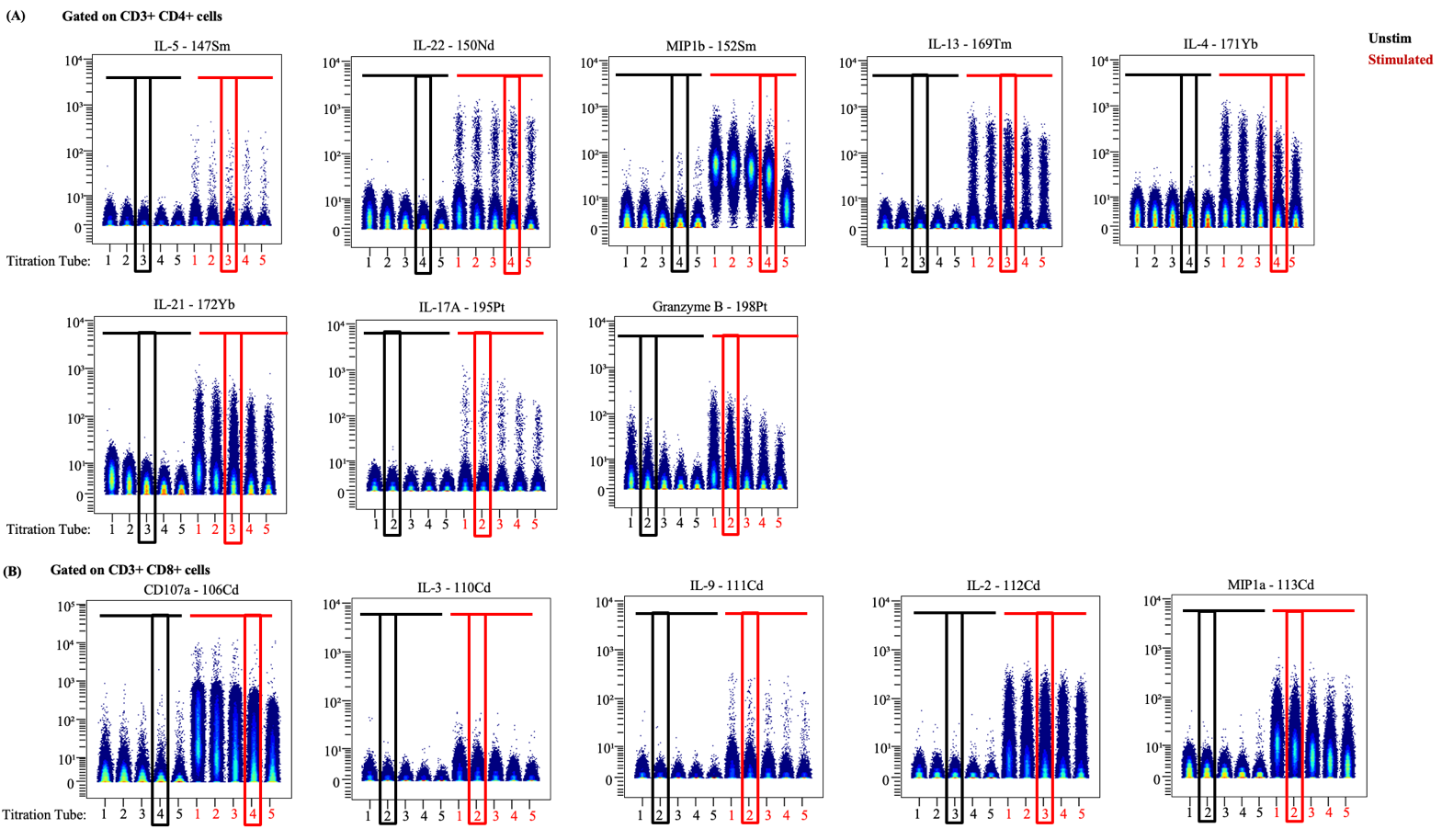

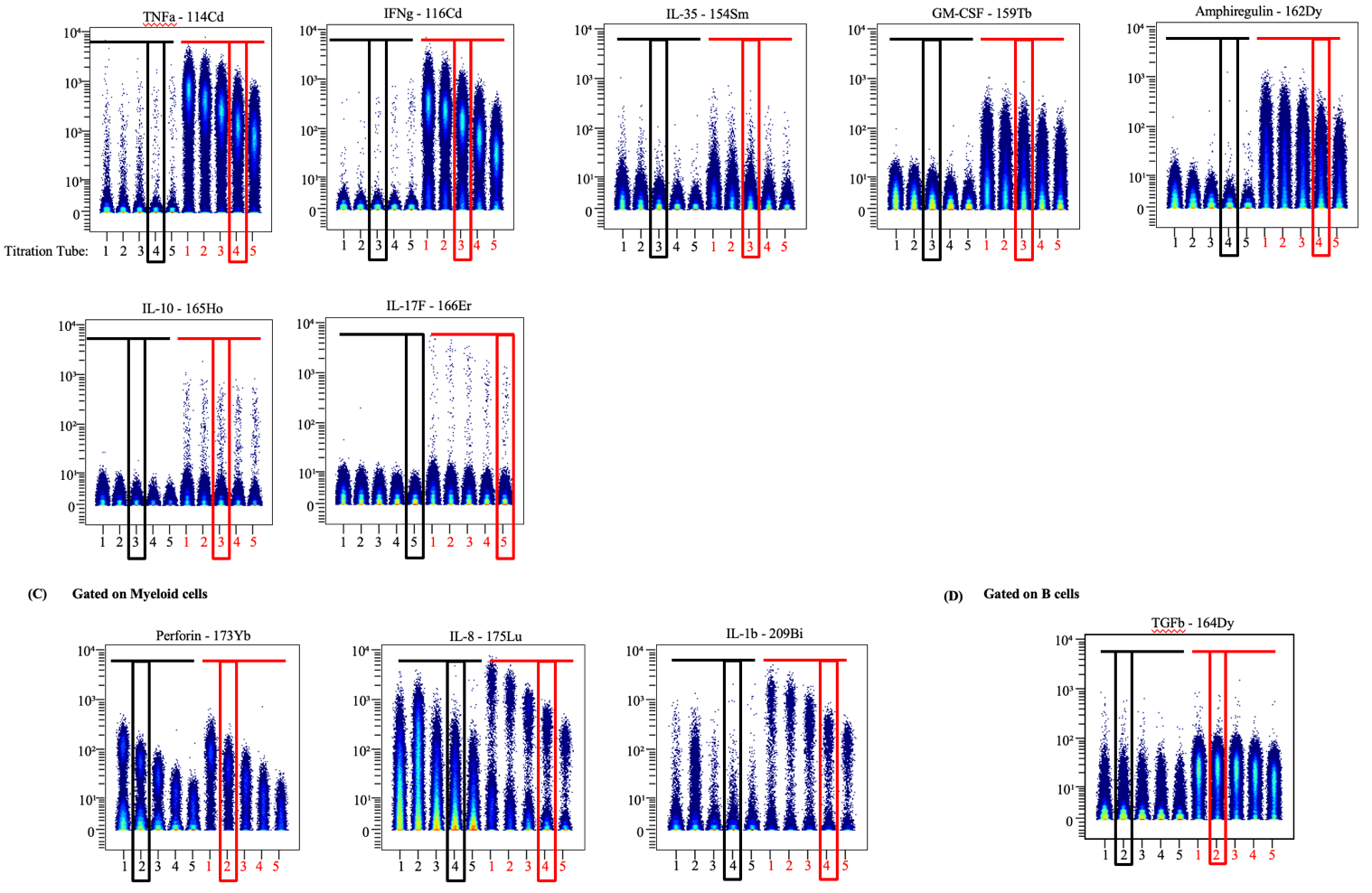
**Supplemental Figure 4. Intracellular marker titrations.** All antibodies were titrated on PBMCs from healthy human donors under both unstimulated and stimulated conditions. Black boxes indicate the selected optimal titration within the unstimulated samples (T1-5), and the red boxes indicate the same selection within the stimulated samples (T6-10).
